# Increased vulnerability to noise exposure of low spontaneous rate type 1C spiral ganglion neuron synapses with inner hair cells (Pre-Print)

**DOI:** 10.1101/2025.05.19.654865

**Authors:** Daniel O. J. Reijntjes, Kali Burke, Srijita Paul, Ulrich Mueller, Elisabeth Glowatzki, Amanda M. Lauer

## Abstract

The inner hair cells (IHCs) in the inner ear form synapses with auditory nerve fibers (ANFs) that send sound signals to the brain. ANFs have been grouped by their level of spontaneous firing rates (SRs) into high-, medium-, and low-SR ANFs. Based on their molecular profiles evaluated by RNAseq experiments, ANFs have been divided into three groups (1A, 1B, and 1C) that likely correspond to high-, medium-, and low-SR ANFs, respectively. In guinea pigs, the synapses between IHCs and low-SR ANFs have been shown to be more vulnerable to noise exposure compared to other ANF subtypes, but not in a study performed in CBA/CaJ mice, questioning if these results can be generalized. Here, an *LYPD1* reporter mouse model on a C57Bl/6J background with specifically labeled group 1C, low-SR ANFs was used to examine whether *LYPD1* positive ANF synapses are more vulnerable to noise exposure. Six-week-old mice were exposed to an 8-16 kHz octave band noise presented at 100 dBA for 2 hours. One week later, cochlear tissue was harvested to quantify ANF synapses and compare the percentage of *LYPD1* positive ANF synapses in noise-exposed and unexposed animals. Auditory brainstem response measurements were performed to assess hearing function after noise exposure. The number of all ANF synapses and the percentage of *LYPD1-*positive ANF synapses were reduced following noise exposure, concurrent with increased ABR thresholds and decreased ABR wave 1 amplitudes. The reduction in the percentage of *LYPD1-*positive ANF synapses specifically indicates greater vulnerability of *LYPD1* positive ANF synapses to noise exposure compared to other ANFs in C57Bl/6J mice.

## 1. Introduction

Sound information is encoded into electrical signals by inner hair cells (IHCs) that form synaptic connections with 10-20 auditory nerve fibers (ANFs) per IHC in the mammalian cochlea (Meyer and Moser 2010). Each IHC forms synaptic connections to a complement of ANFs with different physiological properties which likely serves to increase the range of sound intensities an individual IHC can encode (Liberman 1978). These IHC synapses have a specialized presynaptic ribbon structure that contains the ribeye protein (Schmitz et al. 2006) and the ribbon structure likely facilitates rapid and sustained release of glutamate-filled vesicles (Matthews and Fuchs 2010). The released glutamate activates glutamate receptors which are clustered at the synaptic cleft in the postsynaptic ANFs alongside other necessary synaptic machinery (Verpelli et al. 2012). These afferent synapses are spatially segregated at the IHC base, whereby the ANFs with low-spontaneous rates (low-SR) and high stimulation thresholds preferentially form synapses on the modiolar side of IHCs, whereas ANFs with high-spontaneous rates (high-SR) and low stimulation thresholds preferentially form synapses on the pillar side of IHCs, and ANFs with medium spontaneous rates and thresholds in between (Merchan-Perez and Liberman 1996; Siebald et al. 2023). The spontaneous rates correlate with subtle differences in the diameter of the ANFs and the size of synaptic components, including ribbon size (Liberman, Wang, and Liberman 2011; Stamataki et al. 2006). The underlying mechanisms that set the differences in the physiological properties of these neurons are not completely known but likely include pre-synaptic (Niwa et al. 2021; Özçete and Moser 2021) and postsynaptic (Rutherford et al. 2023) mechanisms.

Several studies suggest that the synapses of ANFs with low-SR are more vulnerable to damage than high-SR ANFs as seen in the relatively larger loss of such ANFs in single unit recordings following noise exposure in albino guinea pigs (Furman, Kujawa, and Liberman 2013; Song et al. 2016), cochlear perfusion of kainite in gerbils (Diuba et al. 2025) and during aging in gerbils and mice (Schmiedt 1996; Wang, Lin, and Xie 2023). However, such a specific loss of low-SR fibers is not so obvious in noise-exposed CBA/CaJ mice (Suthakar and Liberman 2021). To clarify this discrepancy in the apparent vulnerability of low-SR ANFs in mice compared to other species, the vulnerability of low-SR ANFs to noise exposure was re-examined in this study in a different mouse strain. Such an increased vulnerability has significant implications for auditory processing because SR is correlated to ANF properties such as threshold, latency, adaptation and phase locking in response to sound exposure, and loss of such fibers would therefore significantly affect sound coding in the auditory nerve (Costalupes, Young, and Gibson 1984; Joris and Yin 1992; Rhode and Smith 1985). However, ANFs may show plasticity in their response properties following noise exposure depending on the species examined, making it difficult to predict how loss of the low-SR ANFs affects hearing across species (Diuba et al. 2025; Suthakar and Liberman 2021).

The physiological subgroups of ANFs have been proposed to correlate with differences in the genetic make-up of the corresponding spiral ganglion cell bodies as determined by single cell RNA sequencing techniques (Petitpré et al. 2018; Shrestha et al. 2018; Sun et al. 2018). Using clustering analysis on differences in gene expression, these studies suggest that the spiral ganglion neurons’ genetic profiles separate these neurons into at least three groups that seemingly match low-SR, medium-SR and high-SR ANFs, whereby type 1C spiral ganglions would correspond to low-SR ANFs forming synapses on the modiolar side of IHCs, type 1A spiral ganglion neurons would correspond to high-SR ANFs forming synapses on the pillar side of IHCs, and type 1B ANFs would correspond to the medium-SR ANFs forming synapses closer to the basal pole of the IHCs. A genetic mouse model (*Lypd1-CreERT2*^+*/Cre*^) was previously developed where *Lypd1* positive neurons, found specifically in type IC spiral ganglion neurons, were examined to determine that type IC spiral ganglion neurons indeed form synaptic connections on the modiolar side of IHCs and have low-SRs (Siebald et al. 2023).

The approach of the study here to reassess the noise vulnerability of low-SR ANFs in mice, was to use the reporter mouse model with labeled low-SR fibers of the molecularly defined 1C SGN subgroup. Low-SR ANFs were genetically labeled by crossing of *LYPD1-CreERT2*^+*/Cre*^ mice with the *Ai14*^*flox/flox*^ line that expressed a Cre-inducible td-tomato reporter (Madisen et al. 2010; Siebald et al. 2023). *Lypd1-CreERT2*^+*/Cre*^*;Ai14*^+*/flox*^ mice were analyzed for audiometric function using ABR measurements before and after exposure to broadband noise previously shown to induce synaptopathy (Burke et al. 2022; Kujawa and Liberman 2009). After noise exposure, the total number of surviving synapses overlapping with *Lypd1-CreERT2*^+*/Cre*^*;Ai14*^+*/flox*^ positive and negative ANFs were quantified using immunohistochemical analysis.

Noise exposure significantly reduced the number of all IHC/ANF synapses in the high frequency region of the cochlea (32 kHz) measured one week after noise exposure. similar to a study where the averaged number of all IHC/ANF synapses along cochlear apex to base locations was found to be reduced 4-7 days after noise exposure in C57Bl/6J mice (Shi et al. 2015). In addition, the percentage of *Lypd1-CreERT2*^+*/Cre*^*;Ai14*^+*/flox*^ positive to all ANFs was reduced after noise exposure, suggesting an increased vulnerability of low-SR synapses (1C-type) to noise exposure. These data show that, similar to guinea pig (Furman, Kujawa, and Liberman 2013; Song et al. 2016), *Lypd1-CreERT2*^+*/Cre*^*;Ai14*^+*/flox*^ mice on a C57Bl/6J background show an increased vulnerability of low-SR ANFs to noise exposure compared to other ANF subgroups.

## 2. Methods

### 2.1. Animals

Mice used in these experiments were a cross between the *Lypd1-CreERT2* mouse line generated by the Müller lab (Siebald et al. 2023) and the *Ai14* Cre-reporter mouse strain (RRID:IMSR_JAX:007914), both on a C57Bl/6J background. When these mice are crossed, both the native *Lypd1* gene and the *Ai14* gene are transcribed in *Lypd1*^+^ cells. Breeding of *Lypd1-CreERT2*^+*/Cre*^;*Ai14*^*flox/flox*^ animals with *Lypd1-CreERT2*^+*/Cre*^;*Ai14*^*flox/flox*^ animals resulted in offspring either positive or negative for *Lypd1-Cre ERT2*. Genotyping for *Lypd1-CreERT2* was performed using the following primers generated by Riley Bottom in the Müller lab and allowed for assessment of heterozygous or homozygous expression of *Lypd1-CreERT2*.

Primer 1: LYPD1CreERT2_Common FWD CACATATGCTAACCCCACTTCTC

Primer 2: LYPD1CreERT2_Mut Rev GTACGGTCAGTAAATTGGACATTGAG

Primer 3: LYPD1CreERT2_WT Rev GTCTGGACTTTCTGCTCCATG

Twenty-eight *Lypd1-CreERT2*^+^*;Ai14*^*flox/flox*^ mice (15 female, 13 male) were noise-exposed at P43 and euthanized for tissue collection at P50. Animals were randomly assigned to either the “Unexposed” or “Noise exposed” group. Three animals died during the course of the experiments, as a result of tamoxifen injection, meaning 25 animals were analyzed for histology (13 unexposed/12 noise-exposed). Two cochleae were excluded from the analysis due to poor-quality dissections resulting in 48 cochleae included for histological analysis. Animals were socially housed in a low-noise vivarium (Wu et al. 2020), had continuous access to food and water, and were maintained in a 12/12 h light/dark cycle. All experiments were performed in accordance with the protocols approved by the Johns Hopkins University Animal Care and Use Committee and the Guide for the Care and Use of Laboratory Animals.

To control for any differences in baseline audiometric function between animals that were positive or negative for *Lypd1-CreERT2*^+^*;Ai14*^*flox/flox*^, nine animals negative for *Lypd1-CreERT2*^+*/Cre*^;*Ai14*^*flox/flox*^ were used to compare the baseline hearing function to the animals positive for *Lypd1-CreERT2*^+^*;Ai14*^*flox/flox*^ and no difference was found (supplementary Figure 1). These animals were not included in the assessment of noise exposure.

It should be noted here that preliminary ABR testing performed on a cohort of *Lypd1-CreERT2*^+*/Cre*^ animals crossed with the *Ai9* reporter mouse strain (RRID:IMSR_JAX:007909) indicated that these mice already showed an increased ABR thresholds at baseline compared to *Lypd1-CreERT2*^+*/*+^ controls not expressing *CreERT2* (data not shown), necessitating the switch to *Ai14* mice that had normal ABR thresholds.

### 2.2. Auditory brainstem response measurements

Auditory brainstem response (ABR) testing procedures were similar to those previously described (Burke et al. 2024; Capshaw et al. 2022; Mondul et al. 2024). Briefly, ABR testing was conducted in a sound-attenuating chamber (Industrial Acoustics Company, Bronx, NY; 59 × 74 × 60 cm) lined with 4 cm thick acoustic foam (Pinta Acoustic, Minneapolis, MN). Mice were anesthetized with an intraperitoneal (i.p.) injection of 100 mg/kg ketamine and 20 mg/kg xylazine and placed on a heating pad to maintain a temperature of 37 °C. Subdermal stainless steel needle electrodes (Disposable Horizon, 13 mm needle, Rochester Med, Coral Springs, FL) were placed on the vertex (active), bulla (reference), and hind limb (ground). One ear was tested per subject (random and counterbalanced selection, consistent across time points).

ABRs were recorded in response to broadband clicks (100 μs) and tone bursts (8, 12, 16, 24, 32, 42 kHz; 5 ms duration, 0.5 ms rise/fall) at a rate of 21/s for a total of 512 presentations with alternating stimulus polarities. Stimuli were created in SigGen software (Tucker-Davis Technologies; TDT, Alachua, FL) and generated by an RZ6 multi-I/O processor (TDT). Stimuli were played from a free field speaker (MF1, TDT) located 10 cm from the mouse’s pinna at 0° azimuth. Stimuli were calibrated with a 0.25 in. free-field microphone (PCB Piezotronics, Depew, NY, model 378C01) placed at the location of the mouse’s ear. Stimulus levels (decibel; dB) ranged from 90 to 10 dB SPL, decreasing in 10 dB steps. ABR signals were acquired with a Medusa4Z preamplifier (12 kHz sampling rate) and filtered from 300–3000 Hz with an additional band-reject filter at 60 Hz. Post hoc filters from 300– 3000 Hz with steeper cutoff slopes were also applied for additional smoothing. ABRs were measured at baseline and 7 days after noise or sham exposure and tamoxifen injection. Once the ABR was complete, mice were removed from the chamber and placed on a heating pad to recover before being returned to their home cages.

ABR waveforms were analyzed offline by a blinded observer and then verified/corrected by a second blinded observer. Inter-rater reliability was >0.85 for ABR thresholds. ABR threshold was defined as the intermediate intensity between the lowest sound level to evoke a response and when that response was no longer visible above the noise floor (e.g., waves present at 50 dB waves absent at 40 dB, threshold = 45 dB). Peak-to-trough amplitudes and peak latencies were derived for ABR Wave I using a semi-automated ABR wave analysis software previously described in (Burke, Burke, and Lauer 2023) (https://github.com/mattbke63/Auditory-Brainstem-Response-Waveform-Analysis). Due to our hypotheses relating to the status of the auditory nerve, we focused our analyses on only wave 1. ABR thresholds, wave 1 amplitudes, and wave 1 latencies were converted to shifts from baseline to normalize across animals and to reduce the complexity of our statistical models.

### 2.3. Noise exposures and tamoxifen injections

Noise exposures were conducted in awake animals at P43 in a sound-attenuating booth (Industrial Acoustics Company; 37 × 53 × 33 cm) lined with 4 cm thick acoustic foam (Pinta Acoustic, Minneapolis, MN). The noise exposure was 8-16 kHz octave band 100 dBA for 2 hours, and the sham exposure mice were placed in the same chamber with no sound turned on (see Burke et al., 2022 for results from a similar exposure in CBA/CaJ mice). The noise-exposure was controlled by a Hewlett Packard xw4400 Workstation, and mice were restrained in a small 5.5 x 5 x 10.5 cm chicken wire holder 11 cm underneath a PRO Master USA 2” Titanium Super Tweeter (Model TW-47) presenting the noise. The noise was amplified using a CrownD-75A amplifier. The sound intensity was calibrated before and after the exposure using a Larson Davis LxT2 sound level meter with a ½ inch microphone. Immediately following exposures, all mice, including the *CreERT2* negative animals, were given an i.p. injection of 0.2 ml of Tamoxifen (Sigma #T5648) dissolved in corn oil by sonication (Sigma #C8267) at a concentration of 10 mg/ml to express Cre-Lox recombination before being returned to their home cage.

### 2.4. Histology

On the same day of the final ABR recording, mice were anesthetized using isoflurane inhalation (Forane, Baxter Healthcare Corporation, USA) and decapitated after absence of foot withdrawal reflexes. The cochleas were removed from the skull and dissected in ice cold PBS. Two holes exposing the apical and basal turns were chipped into the cochlear bone using Dumont #5 forceps (fine science tools). Cochleas from both ears were then incubated in 4% paraformaldehyde (PFA, Electron microscopy sciences 15714) on ice for 1 hour. The entire epithelial coil containing the organ of Corti was then excised in ice cold PBS, using a custom tool to detach the lateral wall from the cochlear bone during the dissection leaving as much of the lateral wall intact as possible, and placed in blocking buffer (5% normal goat serum, 0.6% triton-X and 0.5% saponin in PBS) for at least 1 hour at 4°C. Organs of Corti were incubated in the lids of 1.5 ml microcentrifuge tubes in primary antibody overnight at room temperature (RT). Two initial litters were processed with the following primary antibodies: mouse IgG 1 anti CTBP2 (BD Biosciences 612044), rabbit anti GluA2/3 (Milipore AB1506), and chicken anti neurofilament (Milipore AB5539) using the endogenous Ai14 expression of *LYPD1* positive neurons to count *LYPD1* positive fibers. The endogenous Ai14 expression was quite variable across samples and, therefore, a further two litters were processed with the following antibodies: mouse IgG 1 anti CTBP2 (BD Biosciences 612044), chicken anti neurofilament (Milipore AB5539), rabbit anti red-fluorescent protein (Rockland 600-401-379) and mouse IgG 2a anti post-synaptic-density protein 95 (PSD95, Neuromab 75-028). All primary antibodies were used at a dilution of 1:600. Organs of Corti were rinsed three times in PBT (PBS with 0.6% triton-X). Organs of Corti were then incubated in secondary antibody at RT at a dilution of 1:1200. For the initial set of experiments the following secondary antibodies were used: alexa 405 anti-chicken (Biotium custom antibody), alexa 488 anti-rabbit (invitrogen A11070), alexa 631 anti-mouse IgG 1 (invitrogen A21126). For the second set of experiments the following secondary antibodies were used: 405 anti-chicken (Biotium custom antibody), alexa 488 anti-mouse IgG 1 (invitrogen A21121), alexa 568 anti-rabbit (invitrogen A11036), alexa 647 anti-mouse IgG 2a (invitrogen A21241). Organs of Corti were rinsed a further three times in PBT and incubated in 125 mM EDTA (Quality Biological 351-027-101) for 15 minutes and rinsed a further three times in PBT before being stored in PBS. Organs of Corti were then cut into a basal, middle and apical turns using fine scissors (fine science tools 14060-10). The lateral wall was removed using a scalpel blade (Bard-parker size 15, 371215). Organs of Corti were then mounted on slides in vectashield mounting medium (Vectorlabs H-1000-10).

### 2.5. Imaging

Low-magnification images of each cochlear piece were taken using a Nikon Eclipse 80i light microscope. Place-frequency maps were created for each cochlea using the ImageJ plugin provided by the Eaton-Peabody-laboratories https://masseyeandear.org/research/otolaryngology/eaton-peabody-laboratories/histology-core) based on a published place frequency map by (Müller et al. 2005) with the caveat that the place frequency map changes following noise exposure (Müller and Smolders 2005). Using these maps, a total of 78 confocal images were taken of the 8, 16 and 32 kHz regions using a Nikon A1 HD25 (with 25 mm field of view) confocal microscope with a 60x = OIL Plan APO NA 1.40 (working distance 0.13 mm) objective. The pinhole size was 0.7 airy unit. The zoom was 200% to capture 15-20 inner hair cells (IHCs) per scan. After acquisition, the Nikon denoising algorithm was applied to each scan. Scans were done sequentially with the 405 and 568 and 488 and 647 channels acquired simultaneously without averaging.

### 2.6. Image analysis

The number of synapses in the ‘noise-exposed’ group or ‘unexposed’ group were counted using the IMARIS software version 11 (Oxford instruments). A ‘synapse’ was defined as to contain a presynaptic CTBP2 punctum juxtaposed to either a GluA2/3 or PSD95 postsynaptic punctum. Using the automated “spots” function, all CTBP2 puncta were counted and checked by eye. A second spots function was then applied to either GluA2/3 or PSD95 to count the number of postsynaptic puncta. All puncta not associated with a CTBP2 punctum were excluded after checking by eye. This second analysis was used as quantification for the number of synapses. Eight samples were excluded where the amount of noise in the channel containing the PSD95 signal was too high to accurately count the postsynaptic endings leaving 44 samples. The number of *LYPD1* positive synapses was counted for both the *Lypd1-CreERT2;Ai14* signal or the RFP signal by hand. ‘*Lypd1*^+^ synapses’ were counted, whenever *Lypd1*^+^ auditory nerve fibers overlapped with the postsynaptic marker of a synapse. For 26 out of 78 scans from the samples where the endogenous *Lypd1-CreERT2;Ai14* labeling was used to assess whether neurons were *LYPD1* positive, the signal was too weak to confidently determine whether neurons were positive for *LYPD1* or not and these were excluded. All histological analysis was performed blinded to experimental group.

### 2.7. Data analysis

All data were analyzed using Linear mixed effect models with animal ID as random fixed effect. For each test, a step-down approach was used to find a minimally-adequate-model examining the relationship between the dependent variable (either; threshold shift, wave 1 amplitude shift, wave 1 latency shift, or synapse count) and several predictors; exposure-group (unexposed or noise-exposed), stimulus-frequency (8, 12, 16, 24, 32, or 42 kHz, this factor not included for click analyses), and stimulus-intensity (50-90 dB, suprathreshold for all exposure groups and time points). Sex was included as a factor in the model for threshold shifts but was excluded from all other models to avoid potential four-way interactions. Post hoc analyses were performed by comparing the estimated marginal means of the models by manually setting contrasts to compare unexposed versus noise-exposed groups for each frequency corrected for by Bonferroni correction. For the synapse count analysis, cochleas from the “noise-exposed” group were excluded if the threshold shift seven days after noise was <= 5dB-SPL to ensure the assessed ears had at least a 10 dB-SPL threshold shift leading to data from 7 animals to be excluded.

All analyses were performed, and all graphs were made in R (R core team 2021) using the following packages: readxl, lme4, lmerTest, emmeans, ggplot2, ggpubr, ggbeeswarm, Matrix, car, dplyr, Rmisc, writexl.

## 3. Results

To examine whether *Lypd1* positive ANFs, representing a low SR, type 1C ANF subgroup, are more vulnerable to damage from noise exposure compared to other ANF subtypes, a previously created *Lypd1* reporter mouse line (*Lypd1-CreERT2*^+*/Cre*^*;Ai14*^*flox/flox*^ mice) on a C57Bl/6J background was used (Siebald et al. 2023) and these animals were exposed to noise levels previously shown to induce a permanent threshold shift in CBA/CaJ mice (Burke et al. 2022). Littermate mice were exposed at P43 to either a 100 dBA 8-16 kHz octave band noise for 2 hours in a noise booth (noise exposure group) or, were kept in the same booth for 2 hours with the noise off (unexposed group). Baseline audiometric function was measured 3 days before noise exposure (baseline -3d) at P40, and vulnerability to noise was assessed seven days after noise exposure (7d) for both groups using auditory brainstem response (ABR) measurements to quantify changes to both auditory thresholds and suprathreshold wave 1 responses. Animals were sacrificed following the ABR after noise exposure and cochlear tissue was processed for histological examination of the number of *Lypd1* positive and negative synapses. To assess relative vulnerability of the *Lypd1* positive ANFs, the percentage of *Lypd1* positive synapses of all synapses was calculated and compared between noise-exposed and unexposed animals. All experiments were performed in *Lypd1-CreERT2*^+*/Cre*^*;Ai14*^*flox/flox*^ (positive) mice.

### 3.1. Threshold shifts after noise exposure

Seven days after noise exposure, ABR thresholds increased by 8 - 24 dB in the ‘noise exposure group 7d’ compared to the ‘noise exposure baseline group -3d’ (Figure 1A, model estimated data is shown in supplementary Table 1). In the same time frame, ABR thresholds in the ‘unexposed group 7d’ decreased by 5 - 15 dB compared to the ‘unexposed baseline group -3d’. Averaged ABR traces for 70 dB SPL click stimuli show a diminished response in the ‘noise exposure group 7d’ compared to the ‘unexposed group 7d’ (Figure 1B). Noise-induced threshold shifts in the unexposed and noise-exposed groups across all measured frequencies were statistically evaluated using a linear mixed effects model. The model included stimulus frequency, exposure group and sex as predictors with no interactions included as the interactions did not contribute significantly to the model. All predictors significantly affected threshold shift (exposure group F(1,28) = 31, p < 0.0001; stimulus frequency F(6,364) = 2.22, p = 0.04; sex F(1,28) = 4.5, p = 0.044). Post hoc comparisons of threshold shifts between unexposed and noise-exposed groups were performed for each stimulus frequency split by sex. For both the males and females, significant differences in threshold shift were observed for all stimuli (Figure 1C-D). These results demonstrate that noise exposure induced a threshold shift lasting up to at least seven days after noise exposure for both males and females. This suggests that significant damage was induced to auditory structures.

**Figure 1.**
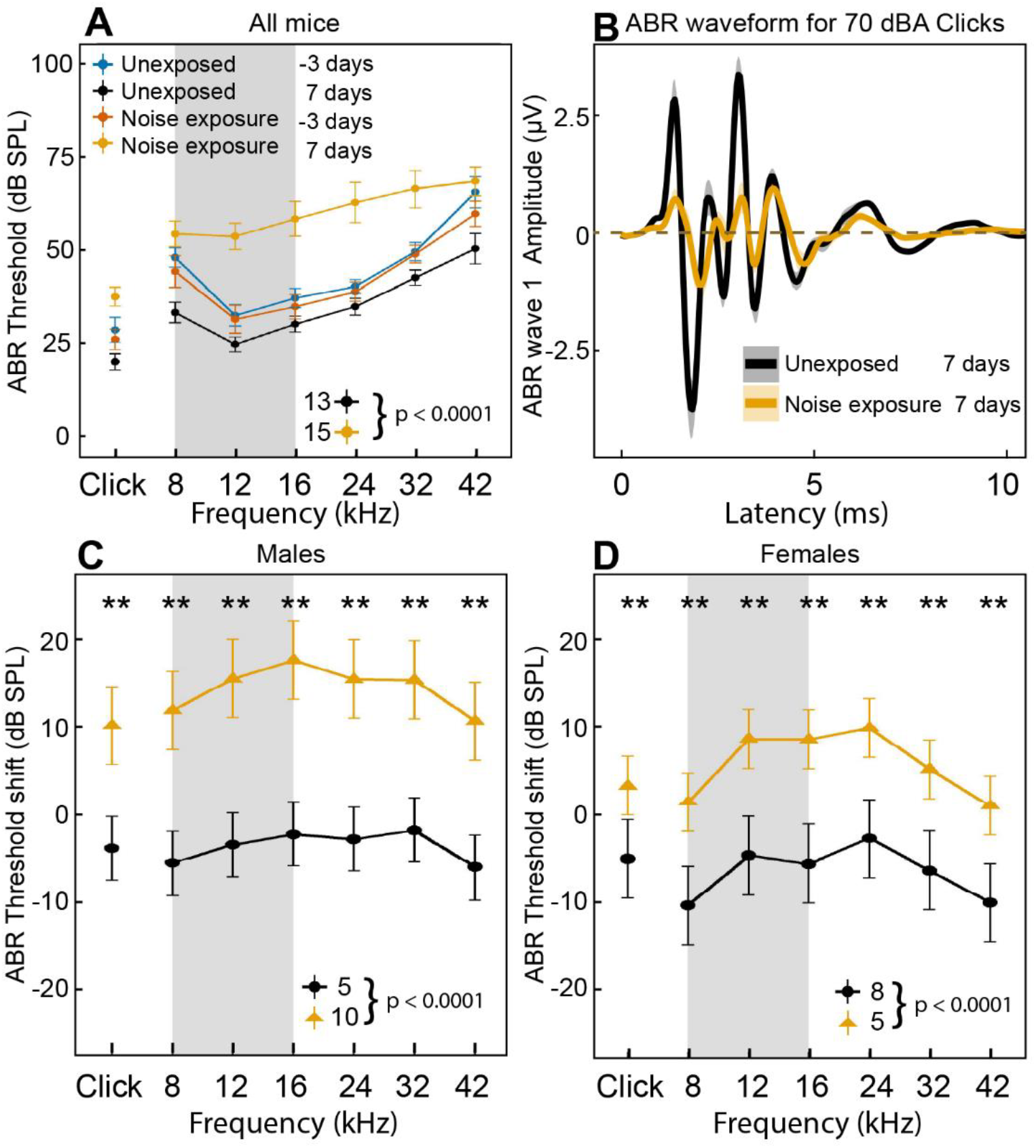
Thresholds and threshold shifts before and after noise exposure, in noise-exposed and unexposed animals. Noise exposure consisted of 2 hours of exposure to a 100 dBA 8-16 kHz octave band noise, as indicated by the grey bars in A, C-D. ABRs were measured 3 days before (-3 days) and 7 days after noise exposure (7 days) in unexposed and noise-exposed animals. Error bars show S.E.M.s (A, C, D) and the shaded area around the waveform represents S.E.M. (B). **A)** Baseline ABR thresholds (-3 days) were similar for both the ‘unexposed group baseline -3d’ and the ‘noise exposure group baseline -3d’. At 7 days after noise exposure, ABR thresholds improved slightly for the ‘unexposed group 7d’, whereas ABR thresholds of the ‘noise exposure group 7d’ worsened compared to all other tested groups. **B)** Grand average ABR traces for 70 dB SPA Click stimuli show a decreased response in the ‘noise exposure group 7d’ compared to the ‘unexposed group 7d’. **C-D)** Sex significantly affected threshold shift; therefore, threshold shift is shown separately for males and females. For both sexes, there was a significant effect of noise exposure across all frequencies linear mixed effects model *p<0.0001, error bars indicate SEM. P-values shown in the panels represent the model statistics. The asterisks indicate the results of the post hoc analysis where ** = p < 0.001. Post hoc analysis showed a significant difference at all frequencies. The N in each group is shown in each panel.

### 3.2. Auditory brainstem response wave 1 suprathreshold responses

To assess suprathreshold responses, and particularly the effect of noise exposure on ABR wave 1 suggestive of damage to auditory nerve fibers, both, amplitudes and latencies of the ABR wave 1 for click and tone stimuli were assessed. Because of the order of magnitude difference in amplitude in the responses to click stimuli, ABR wave 1 responses to click stimuli were statistically analyzed separately from tone stimuli.

### 3.3. Wave 1 amplitude shifts for click stimuli

To account for relative changes for both the exposed and unexposed groups, amplitude shift was calculated for all mice and compared between exposed and unexposed animals. Amplitude shift was calculated by subtracting the amplitude of wave 1 seven days after noise exposure from the amplitude of wave 1 at baseline (-3d) for each animal in the ‘noise exposure group 7d’ and in the ‘unexposed group 7d’ and compared between unexposed and noise-exposed animals using a linear mixed effects model. The minimally adequate model included exposure group as a significant predictor of wave 1 amplitude shift, meaning that wave 1 amplitudes were significantly smaller in the ‘noise exposure group 7d’ (Figure 2A and supplementary table 2, F(1,32) = 11.6, p = 0.002).

**Figure 2.**
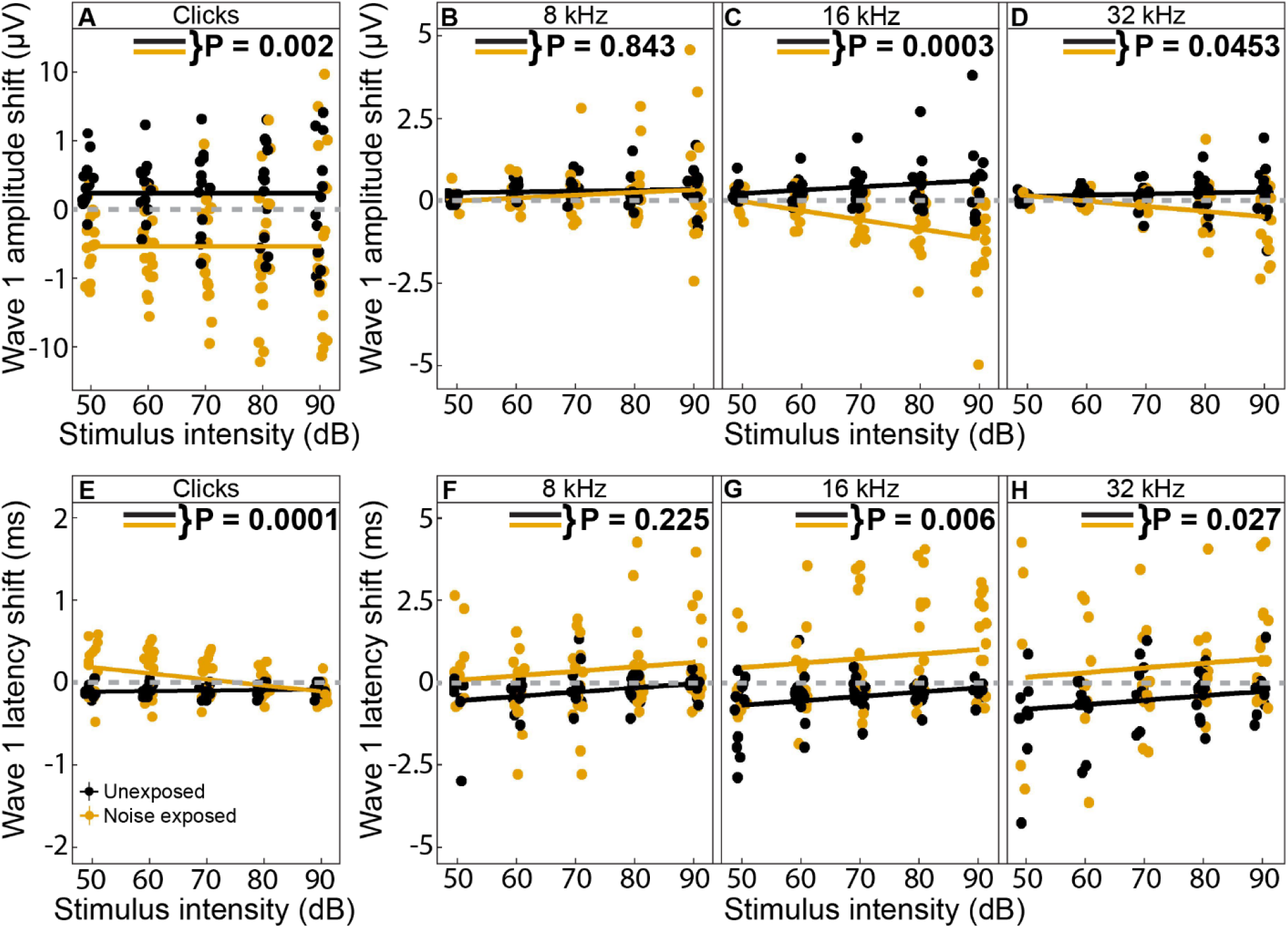
ABR Wave 1 amplitude and latency shifts for click, 8 kHz, 16 kHz and 32 kHz stimuli. Wave 1 amplitudes and latency shifts were calculated by subtracting the amplitude or latency of wave 1 seven days after noise exposure from the amplitude or latency of wave 1 at baseline (-3d). ABR wave 1 amplitude or latency shifts were analyzed by linear mixed effects models. **A)** For click stimuli, in the ‘noise exposure group 7d’, the wave 1 amplitude shift was significantly different from the ‘unexposed group 7d’ meaning the wave 1 amplitude was smaller (F(1,32) = 11.6, p = 0.002). **B-D)**. For tone stimuli, there was a significant three-way interaction between exposure group, stimulus intensity, and stimulus frequency, with the ‘noise exposure group 7d’, wave 1 amplitude shift being significantly different compared to the ‘unexposed group 7d’ for tone stimuli; meaning the wave 1 amplitude was smaller, dependent on the specific frequency and stimulus intensity (F(2,355.6) = 8.84, p = 0.0002). A subsequent analysis split by frequency comparing amplitude shift for the ‘noise exposure group 7d’ to the ‘unexposed group 7d’ averaged over stimulus intensity showed a significantly different ABR wave 1 amplitude shift for the ‘noise exposure group 7D’, at 16 kHz and 32 kHz but not at 8 kHz (8 kHz; t(51.8) = 0.57, p = 0. 843, 16 kHz; t(51.4) = 5.7, p = 0.0003, 32 kHz; t(57.7) = 2.09, p = 0.0453). **E)** For click stimuli, there was a significant two-way interaction between exposure group and stimulus intensity with the wave 1 latency shift being significantly different for the ‘noise exposure group 7d’ compared to the ‘unexposed group 7d’ for click stimuli; meaning the latency was longer, dependent on stimulus intensity (F(1,123) = 37.4, p < 0.0001). Averaged over stimulus intensities, the wave 1 latency shift was still significantly different for the ‘noise exposure group 7d’ (F(1,151)= 42.1, p < 0.0001). **F-H)** For tone stimuli, there was a significant interaction between exposure group and stimulus intensity and a significant effect of frequency, with the wave 1 latency shift being significantly different for the ‘noise exposure group 7d’ compared to the ‘unexposed group 7d’ dependent on the stimulus intensity (Type III ANOVA, F(2,355.5)_stimulus-frequency_ = 3.28, p < 0.0001, F(2,355.5)_exposure-group: stimulus-frequency_ = 3.21, p = 0.04). A subsequent analysis split by frequency comparing latency shift for the ‘noise exposure group 7d’, to the ‘unexposed group 7d’ averaged over stimulus intensity showed a significantly different ABR wave 1 latency shift for the ‘noise exposure group 7d’, at 16 kHz and 32 kHz but not at 8 kHz (8 kHz; t(44.0) = -1.82, p = 0. 225, 16 kHz; t(43.6) = -3.32, p = 0.0057, 32 kHz; t(47.0) = -2.72, p = 0.027). P-values in the panels represent individual comparisons between the ‘noise exposure group 7d’ and the ‘unexposed group 7d’ split by frequency and averaged over stimulus intensity.

### 3.4. Wave 1 amplitude shifts for tone stimuli

Similar to the analysis for the clicks, amplitude shift was calculated by subtracting the amplitude of wave 1 seven days after noise exposure from the amplitude of wave 1 at baseline (-3d) and the calculated amplitude shift was compared between the ‘noise-exposure group 7d’ and the ‘unexposed group 7d’ using a linear mixed effects model. The minimally adequate model included a significant three-way interaction between exposure group, stimulus frequency, and stimulus intensity (Type III ANOVA; F(2,355.6) = 8.84, p = 0.0002) (Figure 2B-D). The result can best be explained as a significantly smaller wave I amplitude (indicated by negative slopes in amplitude shift) in the ‘noise exposure group 7d’ compared to the ‘unexposed group 7d’ dependent on both the specific stimulus intensity and the stimulus frequency. To further analyze for which frequencies there was an effect of exposure group on wave 1 amplitude shift, the wave I amplitude shift was compared between ‘noise exposure group 7d’ and ‘unexposed group 7d’ split by stimulus frequency (8, 16, 32 kHz) and averaged over stimulus intensity. No significant difference was found for 8 kHz stimuli (Figure 2B) and a significant difference in ABR wave 1 amplitude shifts for 16 kHz and 32 kHz stimuli (Figure 2C-D, p-values in the panels reflect the frequency specific analysis. Model estimates are provided in supplementary table 2).

### 3.5. Wave 1 latency shift for click stimuli

To account for relative changes for both the exposed and unexposed groups, the shift in latency for all mice was calculated and the shift between exposed and unexposed animals was compared. Latency shift was calculated by subtracting the latency of wave 1 seven days after noise exposure (7d) from the latency of wave 1 at baseline (-3d) and compared between the ‘noise-exposure group 7d’ and the ‘unexposed group 7d’ using a linear mixed effects model. The minimally adequate model included a significant interaction between exposure-group and stimulus-intensity as a significant predictor of wave 1 latency shift, with a significantly slower wave 1 for the ‘noise-exposure group 7d’ compared to the ‘unexposed group 7d’ dependent on stimulus intensity (F(1,123) = 37.4, p < 0.0001, Figure 2E and supplementary table 2).

### 3.6. Wave 1 latency shift for tone stimuli

As for the clicks, the latency shift of wave 1 seven days after noise exposure was calculated by subtracting the wave 1 latency seven days after noise exposure (7d) from the wave 1 latency at baseline (-3d) and compared between the ‘noise-exposure group 7d’ and the ‘unexposed group 7d’ using a linear mixed effects model. The minimally adequate model included an interaction between stimulus-intensity and exposure-group, and stimulus frequency as significant predictors of latency shift. The significant two-way interaction between exposure group and stimulus intensity means that the wave 1 latency is longer for the ‘noise-exposure group 7d’ compared to the ‘unexposed group 7d’ dependent on the specific stimulus intensity. The significant contribution of stimulus frequency to the model suggests that for at least one of the frequencies, the longer wave 1 latency is not significantly different between the ‘noise-exposure group 7d’ compared to the ‘unexposed group 7d’. To further analyze this effect, latency shift was compared between the ‘noise exposure group 7d’ and ‘unexposed group 7d’ split by stimulus frequency (8, 16, 32 kHz) averaged over stimulus intensity. No significant difference for 8 kHz stimuli was found (Figure 2F) and significant differences for 16 kHz and 32 kHz stimuli were found (Figure 2G-H, p-values in the panels reflect the frequency specific analysis. Model estimates are provided in supplementary table 2).

### 3.7. Relative prevalence of *Lypd1*^+^ synapses following noise exposure

To assess whether *Lypd1*^+^ auditory nerve fiber synapses were relatively more affected by noise exposure than *Lypd1*^*-*^ auditory nerve fiber synapses, the number of all synapses and of *Lypd1*^+^ synapses was determined in the ‘noise-exposure group 7d’ and the ‘unexposed group 7d’ using antibody labeling for both, a pre- and postsynaptic marker. Synapses were defined and counted as a presynaptic CTBP2 punctum (labeling the presynaptic ribbon) juxtaposed to a postsynaptic GluA2/3 or PSD95 punctum (labeling postsynaptic glutamate receptors or the postsynaptic density). ‘*Lypd1*^+^ synapses’ were counted whenever *Lypd1*^+^ auditory nerve fibers overlapped with the postsynaptic marker of a synapse.

Exemplar images taken at the 32 kHz frequency region of cochleas from unexposed and noise-exposed animals show labeling of CTBP2 (Green), PSD95 (magenta), and *Lypd1-Ai14* (orange) (Figure 3). ‘unexposed group 7d’ animals showed a full complement of aligned pre- and postsynaptic puncta (Figure 3A), whereas for the ‘noise-exposure group 7d’ animals, various single non-aligned presynaptic and single postsynaptic puncta were detected, including *Lypd1*^+^ ANFs contacting orphan puncta (Figure 3B, arrowheads point to ‘orphan’ ribbons and arrows to orphan PSD95 puncta’). Analysis of the total number of synapses by linear mixed effects model showed a significant interaction between exposure group and frequency (F_2,81.92_ = 24.6, p<0.0001) meaning there were significantly fewer synapses after noise exposure for the ‘noise-exposure group 7d’ compared to the ‘unexposed group 7d’ depending on the frequency. Post hoc analysis showed that there were significantly fewer synapses at the 32 kHz region in the ‘noise-exposure group 7d’ (Figure 3C, t_(117)_ = - 8.4, p = <0.0003,). Similarly, there were significantly fewer *Lypd1*^+^ synapses in the ‘noise-exposure group 7d’ at 32 kHz (Figure 3D) and the percentage of *Lypd1*^+^ synapses was significantly lower at the 32 kHz region in the ‘noise-exposure group 7D’ compared to the ‘unexposed group 7d’ (Figure 3E).

**Figure 3.**
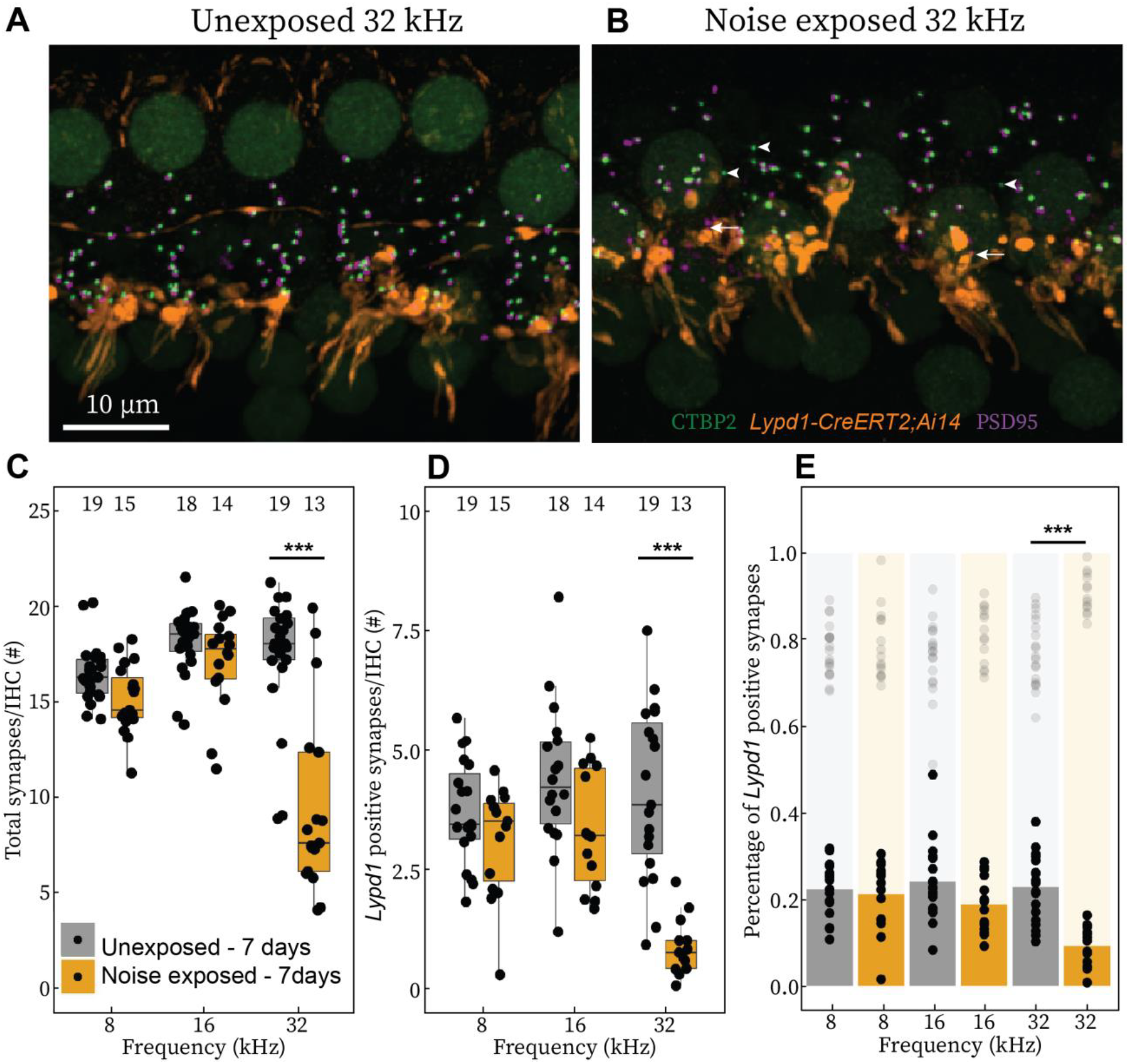
Analysis of total and relative number of *Lypd1*^+^ synapses following noise exposure. A, B). Exemplar images show synapses as pairs of CTBP2 (green) and PSD95 (magenta) and individual *Lypd1*^+^ ANFs (orange) contacting synapses. In the ‘unexposed group 7d’ **(A)**, the majority of synaptic elements are juxtaposed, whereas in the ‘noise-exposure group 7d’ **(B)**, individual pre- (arrowheads) and post-synaptic puncta (arrows) can be seen in addition to a qualitative reduction in the number of *Lypd1*^+^ auditory nerve fibers. **C)** Analysis of the total number of synapses compared between the ‘unexposed group 7d’ and the ‘noise-exposure group 7d’ by linear mixed effects model showed a significant interaction between exposure group and frequency (F_2,81.92_ = 24.6, p<0.0001), meaning there were significantly fewer synapses in the ‘noise-exposure group 7d’ depending on the frequency. Post hoc analysis showed that there were significantly fewer synapses at the 32 kHz region in the ‘noise-exposure group 7d’ (t_(117)_ = -8.4, p = <0.0003). **D)**. Analysis of the total number of *Lypd1*^+^ synapses similarly showed a significant interaction between exposure group and frequency (F_2,64.0.76_ = 12.8, p<0.0001). Subsequent post hoc analysis showed that there were significantly fewer *Lypd1*^+^ synapses in the ‘noise-exposure group 7D’ for the 32 kHz region (t_(89.7)_ = -6.5, p = <0.0003). **E)**. The percentage of *Lypd1*^+^neurons was also compared between ‘unexposed group 7d’ and the ‘noise-exposure group 7d’. The linear mixed effect model showed a significant interaction between exposure group and frequency (F_2,63.7_ = 8.2, p<0.0001). Post hoc analysis showed that the percentage of *Lypd1*^+^ synapses was significantly smaller at the 32 kHz region in the ‘noise-exposure group 7d’ (t_(90.9)_ = -4.9, p = <0.0003, Figure 3E). Boxplots show 25^th^ to 75^th^ percentile and median. Opaque data points represent *LYPD1*^+^ synapses. *LYPD1*^*-*^ data is shown as transparent data points which combined with the *LYPD1*^+^ data points add up to 100%. Significant differences are indicated in the panels (*** corresponds to p-values <0.0001). The numbers above the boxplots indicate the number of samples in that boxplot.

These results confirm that the noise exposure paradigm used in this study significantly reduced the number of cochlear afferent synapses in the highest frequency region examined (32 kHz). In addition, the relative number of *Lypd1*^+^ neurons decreased at the highest frequency region, suggesting that the *Lypd1*^+^ synapses were more vulnerable to noise exposure than *Lypd1*^*-*^ synapses.

## 4. Discussion

In this study the susceptibility to noise exposure of *Lypd1-CreERT2;Ai4*^+^ type 1C ANFs in mice on a C57Bl/6J background was examined. Quantitative evaluation suggests that synapses of type 1C low SR ANFs innervating preferentially the modiolar side of IHCs are more vulnerable to noise exposure compared to synapses formed by other ANF subtypes.

### 4.1. ABR threshold and wave 1 amplitude effects as a result of high-intensity noise exposure

ABR wave 1 threshold increases and suprathreshold wave 1 amplitude decreases following high-intensity noise exposure have been widely reported in multiple strains of laboratory mice (Burke et al. 2022; Kaur et al. 2019; Kujawa and Liberman 2009; Schrode et al. 2018; Sergeyenko et al. 2013; Suthakar and Liberman 2021; Wu, Liberman, and Liberman 2024). Our findings are consistent with the previous literature showing one week after noise exposure an ABR threshold shift accompanied by a reduction in wave 1 amplitude and a reduction of IHC synapses, indicated by a reduction in the number of paired pre- and postsynaptic fluorescent puncta, without the loss of HCs (Kujawa & Liberman, 2009). Similar noise exposure patterns have been most extensively studied in CBA/CaJ mice (e.g. Kujawa and Liberman 2015). CBA/CaJ mice preserve good hearing for most of their lives (Kobrina et al. 2020) whereas C57Bl/6J mice suffer from a mutation in the Cadherin 23 gene (Noben-Trauth et al., 2003), rendering C57Bl/6J mice vulnerable to early onset hearing loss around 3 months of age. Indeed, C57Bl/6J mice differ in their response to noise exposure compared to CBA/CaJ mice (Wu et al., 2024). Despite these differences, the effect of noise exposure on cochlear afferent synapses observed in a C57Bl/6J background in this study was similar to the effect of noise exposure observed in CBA/CaJ mice performed in experiments under similar noise exposure conditions (Burke et al. 2022).

### 4.2. Increased vulnerability of type 1C ANF synapses

Increased vulnerability to noise of *Lypd1*^+^ ANF synapses in mice on a C57Bl/6J background, as reported here in the high frequency region of the cochlea, is consistent with prior work demonstrating preferential loss of low SR ANFs using single unit recordings (citations below), which likely correspond to the type IC ANFs (Siebald et al. 2023). The percentage of low-SR ANFs in guinea pigs is reduced in the total population of single unit recordings after noise exposure (Furman, Kujawa, and Liberman 2013; Song et al. 2016), during aging in gerbils (Heeringa et al. 2023; Schmiedt 1996), and after cochlear kainate perfusion in gerbils (Diuba et al. 2025). Also, histological evidence suggests that type IC ANF somata in the spiral ganglion are more vulnerable to aging in CBA/CaJ mice in the higher frequency region of the cochlea (Wang, Lin, and Xie 2023). However, one study in CBA/CaJ mice with single unit recordings does not report such a preferential loss of low SR fibers (Suthakar and Liberman 2021). In another study, CBA/CaJ mice exhibited almost no recovery of ANFs synapses following noise exposure whereas C57Bl/6J mice did (Wu, Liberman, and Liberman 2024), suggesting that possibly different mechanisms are at work in these two strains.

For the interpretation of the response to noise exposure, time course of damage, possible recovery of IHCs and IHC synapses, and ANF function needs to be considered. For example, in noise-exposed guinea pigs, the median SR of the recorded population of ANFs increased significantly 1 day after noise exposure and recovered roughly after one month (Song et al. 2016). Based on histological measures that assessed the loss of all ANFs regardless of subtype, ANF synapses in mice on a C57Bl/6J background showed some recovery at two weeks post noise exposure (Kim et al. 2019), and in another study after mild noise exposure, ANF synapses recovered after four weeks (Manickam et al. 2023). In the study here, only one timepoint, one week after noise exposure, was investigated which allowed for uncovering low SR ANF vulnerability. However, as synaptic recovery likely was not complete at the one-week time point, it could not be determined if changes were temporary or permanent.

### 4.3. Underlying mechanisms of increased vulnerability of type 1C ANFs to noise exposure

The mechanisms underlying the relatively greater vulnerability of type 1C ANFs remain unclear as well as the functional benefit for type 1C ANFs that would offset this greater vulnerability. The ANFs as a whole show heterogeneity in their physiological properties that appear as gradients along the IHC basal pole. Classical grouping of ANFs based on SR in cat (Merchan-Perez and Liberman 1996) follows a pattern whereby high-SR ANFs contact IHCs on the pillar side and low-, med- and high-SR ANFs are found on the modiolar side. Certain morphological characteristics such as ribbon size (Hua et al. 2021; Liberman, Wang, and Liberman 2011; Stamataki et al. 2006) and ANF diameter (Liberman 1980) are also correlated with these physiological properties and these heterogeneities are reviewed elsewhere (Moser et al. 2023). The type IC ANFs, identified by the *Lypd1-CreERT2;Ai4* mouse line, synapse preferentially on the modiolar side of the IHCs and have some of the lowest SRs recorded, even amongst low-SR ANFs contacting the modiolar side of the IHCs (Siebald et al. 2023). Therefore, it is tempting to assume that type IC ANFs may correspond to the low-SR ANF group found in cat (Merchan-Perez and Liberman 1996).

Pre-synaptic differences between type 1A-C ANFs have not been studied but pre-synaptic differences between pillar and modiolar active zones have been identified and shape how vesicles are released and how glutamate stimulates the postsynaptic ANFs. Pillar presynaptic active zones show tight coupling of the voltage gated calcium channels to release sites, lowering the threshold for vesicular release and likely increasing spontaneous vesicle release events (Özçete and Moser 2021). In contrast, most modiolar presynaptic active zones release vesicles when the IHCs are more depolarized affecting how the post synapse is stimulated. In addition, the maximal presynaptic calcium influx is largest on the modiolar side and correlated to ribbon size (Frank et al. 2009; Ohn et al. 2016). The dynamic range of glutamate release, in terms of IHC depolarization, is smaller for most modiolar presynaptic active zones (Özçete and Moser 2021). Taken together, these findings suggest that the dynamics of glutamate release are different between pillar and modiolar ANFs which likely determines how glutamate is presented to the post-synaptic ANF, perhaps setting up modiolar ANFs, and thus likely the 1C ANFs, to be “overwhelmed” more easily by large volumes of glutamate.

The mode of synapse loss following noise exposure is thought to be glutamate excitotoxicity (Puel et al. 1998), a process whereby overstimulation of the ANF by glutamate leads to intracellular accumulation of Na^+^ and Ca^2+^ with toxic effects (Kostandy 2012). Although the exact mechanisms of excitotoxicity in ANFs are unclear, there is evidence for both Na^+^ and Ca^2+^ influx having toxic effects. Stimulation of ANFs with glutamate causes swelling of the ANF ending (Pujol and Puel 1999) which is likely a result of water entering ANFs as a passive response to the change in intracellular osmolarity due to Na^+^ influx (Kostandy 2012), however, this might be an oversimplified idea (Hellas and Andrew 2021). Furthermore, blocking the calcium permeable subset of glutamate receptors in ANFs (Sebe et al. 2017) reduces glutamate induced and noise induced synapse loss (Hu, Rutherford, and Green 2020; Sebe et al. 2017). Presumably, this effect is mediated by a reduction in the influx of Ca^2+^ into ANF endings. However, blocking part of the AMPA receptors will also reduce the total Na^+^ influx.

Both pathways might affect type IC ANFs more strongly than other ANFs. First, ANF SR is positively correlated to the number of mitochondria found in ANFs (Liberman 1980). As mitochondria can temporarily buffer excess intracellular calcium, fewer mitochondria would make such ANFs more vulnerable to Ca^2+^ influx (Nicholls et al. 2007). Furthermore, long term storage of calcium in mitochondria impairs their function and reduces ATP production which is required for the proper function of ATP-dependent ion transporters (Nicholls et al. 2007) that remove the excess Na^+^ and Ca^2+^ ions. Yet, the exact mechanisms underlying excitotoxicity in the different ANF subgroups remain unclear.

The study here provides clear evidence that type 1C AFNs show higher vulnerability to noise exposure compared to other ANFs, strengthening the need to further investigate underlying mechanisms.

## Acknowledgements

The authors have nothing to disclose. The authors would like to thank Amelie Valles, Sebastian Zamarippa and Michael Nwazue for contributing to preliminary data analysis.

## Supplemental data

**Supplementary Figure 1.**
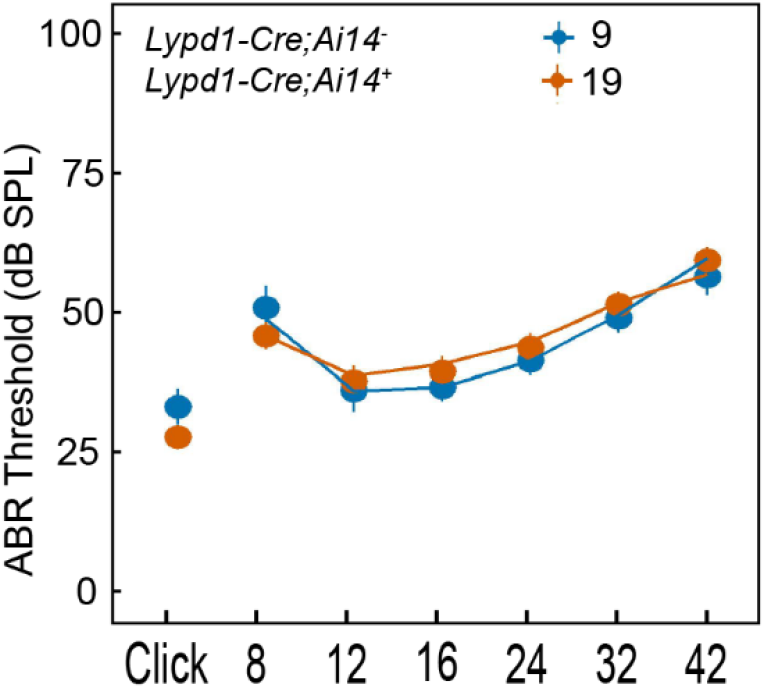
Auditory brainstem response thresholds. Thresholds are shown for both clicks and tones for *Lypd1-CreERT2;Ai14*^+^ (red) and *Lypd1-CreERT2;Ai1*^*-*^ (blue) animals. No differences in ABR threshold were observed at baseline (around P40).

**Supplementary Table 1.** ABR threshold shifts between baseline and 7 days after noise exposure.

| Sex | Stimulus | Unexposed | Unexposed | Noise-exposed | Noise-exposed |
| --- | --- | --- | --- | --- | --- |
|  |  | Mean<br>(dB SPL) | SEM<br>(dB SPL) | Mean<br>(dB SPL) | SEM<br>(dB SPL) |
| Males | Click | -5 | 2.7 | 16 | 5.1 |
|  | 8 kHz | **-11.2** | **3.8** | **24** | **6** |
|  | 12 kHz | **-8.8** | **2.3** | **34** | **4.9** |
|  | 16 kHz | **-3.8** | **2.7** | **34** | **7.8** |
|  | 24 kHz | **-6.3** | **2.5** | **32** | **9.3** |
|  | 32 kHz | **-3.8** | **2.5** | **31** | **6.3** |
|  | 42 kHz | **-12.5** | **4.8** | **22** | **4.9** |
| Females | Click | -14 | 4.0 | 9.0 | 2.8 |
|  | 8 kHz | -20 | 4.0 | 3.0 | 4.0 |
|  | 12 kHz | -6.0 | 3.7 | 16 | 4.7 |
|  | 16 kHz | **-12** | **4.0** | **18** | **5.8** |
|  | 24 kHz | -4.0 | 4.0 | 19.5 | 6.4 |
|  | 32 kHz | -12 | 3.7 | 10.5 | 5.9 |
|  | 42 kHz | -19 | 6.6 | 2.0 | 4.2 |

*Bold numbers indicate significant differences between unexposed and noise-exposed groups after Bonferroni post hoc correction.*

**Supplementary table 2.** Model estimated marginal means.

| Amplitude (μV) | Unexposed E.M.M | Unexposed S.E. | Confidence interval |  | Exposed E.M.M | Exposed S.E. | Confidence interval |  |
| --- | --- | --- | --- | --- | --- | --- | --- | --- |
| Clicks | **1.31** | **0.88** | -0.49 | 3.10 | **-2.45** | **0.73** | -3.94 | -0.97 |
| 8 kHz | 0.28 | 0.15 | -0.03 | 0.58 | 0.16 | 0.13 | -0.10 | 0.42 |
| 16 kHz | **0.42** | **0.15** | 0.12 | 0.73 | **-0.72** | **-0.72** | -0.98 | -0.45 |
| 32 kHz | **0.21** | **0.15** | -0.10 | 0.52 | **-0.22** | **0.14** | -0.50 | 0.06 |
| Latency (ms) |  |  |  |  |  |  |  |  |
| Clicks | **-0.05** | **0.04** | -0.12 | 0.03 | **0.15** | **0.03** | 0.08 | 0.22 |
| 8 kHz | -0.11 | 0.11 | -0.33 | 0.11 | 0.15 | 0.09 | -0.03 | 0.33 |
| 16 kHz | **0.15** | **0.11** | -0.36 | 0.07 | **0.32** | **0.09** | 0.14 | 0.51 |
| 32 kHz | **-0.22** | **0.11** | -0.44 | 0.002 | **0.18** | **0.10** | -0.02 | 0.37 |

*E.M.M = estimated marginal means.*
*S.E. = standard error and represents the estimated standard errors for the estimated marginal means.*
*Bold numbers represent significant differences.*

